# Resolving the origins of membrane phospholipid biosynthesis genes using outgroupfree rooting

**DOI:** 10.1101/312991

**Authors:** Gareth A. Coleman, Richard D. Pancost, Tom A. Williams

## Abstract

One of the key differences between Bacteria and Archaea are their canonical membrane phospholipids, which are synthesized by distinct biosynthetic pathways with non-homologous enzymes. This “lipid divide” has important implications for the early evolution of cells and the type of membrane phospholipids present in the last universal common ancestor (LUCA). One of the main challenges in studies of membrane evolution is that the key biosynthetic genes are ancient and their evolutionary histories are poorly resolved. This poses major challenges for traditional rooting methods because the only available outgroups are distantly related. Here, we address this issue by using the best available substitution models for single gene trees, by expanding our analyses to the diversity of uncultivated prokaryotes recently revealed by environmental genomics, and by using two complementary approaches to rooting that do not depend on outgroups. Consistent with some previous analyses, our rooted gene trees support extensive inter-domain horizontal transfer of membrane phospholipid biosynthetic genes, primarily from Archaea to Bacteria. They also suggest that the capacity to make archaeal-type membrane phospholipids was already present in LUCA.

## Introduction

Archaea and Bacteria form the two primary domains of life (reviewed in Williams et al. 2013). While similarities in their fundamental genetics and biochemistry, and evidence of homology in a near-universally conserved core of genes (Weiss et al. 2016) strongly suggest that Archaea and Bacteria descend from a universal common ancestor (LUCA), they also differ in ways that have important implications for the early evolution of cellular life. These differences include DNA replication (Kelman and Kelman 2014), transcription (Bell and Jackson 1998), DNA packaging (Reeve et al. 1997), and cell wall compositions (Kandler 1995). One striking difference is in the phospholipid composition of the cell membranes (Figure 1), which is particularly important for understanding the origin of cellular life. Canonically, Archaea have isoprenoid chains attached to a glycerol-1-phosphate (G1P) backbone via ether bonds, and can have either membrane spanning or bilayer-forming phospholipids (Lombard et al. 2012a). Most Bacteria, as well as eukaryotes, classically have acyl (fatty acid) chains attached to a glycerol-3-phosphate (G3P) backbone via ester bonds and form bilayers (Lombard et al. 2012a), although a number of exceptions have been documented (Sinninghe Damsté et al. 2002; Weijers et al. 2006; Sinninghe Damsté et al. 2007; Goldfine 2010). These phospholipids are synthesised by non-homologous enzymes via different biosynthetic pathways (Figure 1). This so-called ‘lipid divide’ (Koga 2011) raises some important questions regarding the early evolution of cellular life, including the nature of the membrane phospholipids present in LUCA and the number of times cell membranes have evolved.

**Figure 1.**
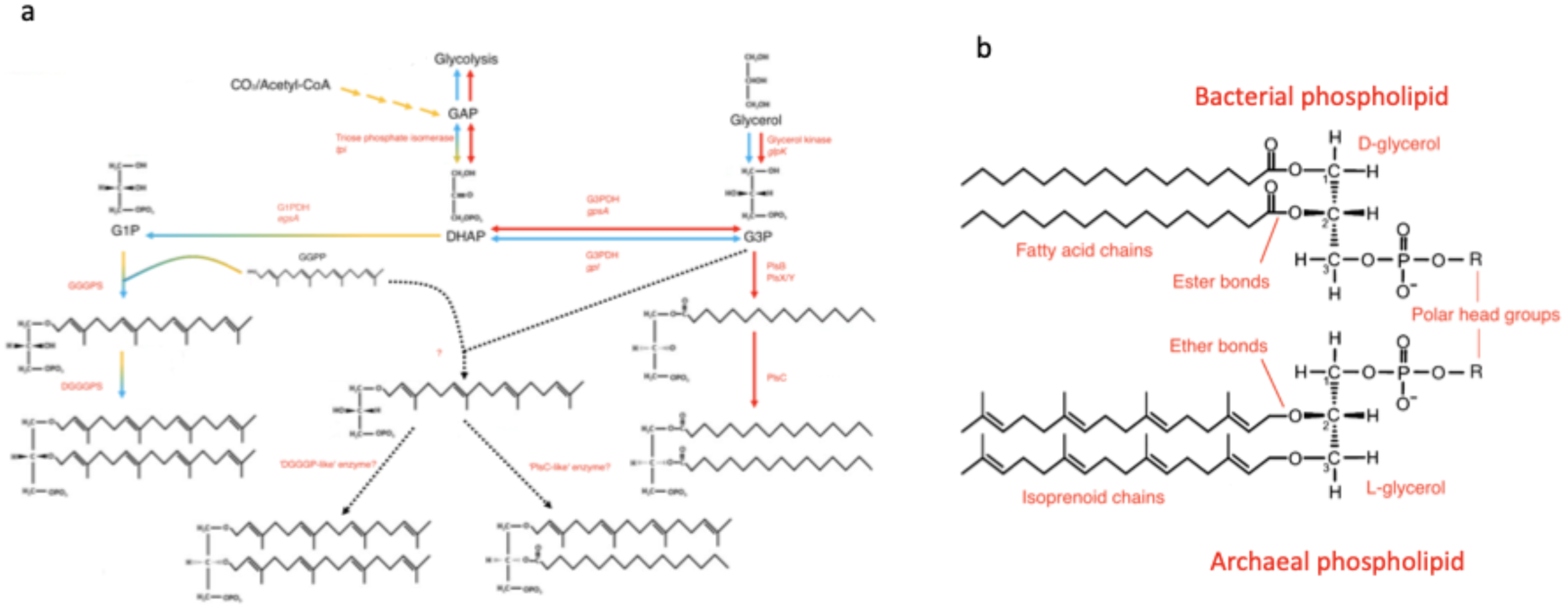
**a)** The canonical ether/ester biosynthetic pathways in Archaea and Bacteria and how they relate to glycerol metabolism. Based on Figure 1 from Villanueva et al (2016). Archaeal pathways in blue and yellow (blue=heterotrophic Archaea, yellow=autotrophic Archaea), bacterial pathway in red. Hypothetical biosynthetic pathway, as suggested by Villanueva et al. (2016), in dashed lines. **b)** Composition of bacterial and archaeal phospholipids. In Archaea, G1P is synthesised from dihydroxyacetone phosphate (DHAP) using the enzymes glycerol-1-phosphate dehydrogenase (G1PDH). The first and second isoprenoid chains (GGGPs) are added by geranylgeranylglyceryl synthase (GGGPS) and digeranylgeranylglyceryl synthase (DGGGPS) respectively. In Bacteria, G3P is synthesised by glycerol-3-phosphate dehydrogenase (G3PDH) from DHAP. There are two forms of this enzyme, encoded by the *gpsA* and *glp* genes respectively. G3P may also be produced from glycerol by glycerol kinase (glpK). In certain Bacteria, such as Gammaproteobacteria, the first fatty-acid chain is added by a version of glycerol-3-phosphate acyltransferase encoded by the *PlsB* gene. Other Bacteria, including most gram positive bacteria, use a system which includes another glycerol-3-phosphate acyltransferase encoded by *PlsY*, in conjunction with an enzyme encoded by *PlsX* (Yao and Rock 2013; Parsons and Rock 2013). The second fatty-acid chain is attached by 1-acylglycerol-3-phosphate O-acyltransferase, encode by *PlsC*.

The observation that phospholipid biosynthesis in Bacteria and Archaea is non-homologous has motivated various hypotheses on the nature of LUCA’s membrane. The likely presence of some genes for lipid biosynthesis (Lombard and Moreira 2011; Lombard et al. 2012a; Koga 2014; Weiss et al. 2016) and, in particular, a membrane-bound ATPase (Sojo et al. 2014; Weiss 2016) in reconstructions of LUCA’s genome implies that LUCA possessed a membrane, although its properties may have been somewhat different to those of modern, ion-tight prokaryote cell membranes (Lombard et al. 2012a; Koga 2014; Sojo et al. 2014). It has also been suggested that LUCA may have had a heterochiral membrane (Wächtershäuser 2003), with later independent transitions to homochirality in Bacteria and Archaea, driven by increased membrane stability. However, the available experimental evidence - including the recent engineering of an *Escherichia coli* cell with a heterochiral membrane (Caforio et al. 2018) - suggests that homochiral membranes are not necessarily more stable than heterochiral ones (Fan et al. 1995; Shimada and Yamagishi 2011; Caforio et al. 2018), requiring some other explanation for the loss of ancestral heterochirality.

Despite the importance of the lipid divide for our understanding of early cellular evolution, membrane phospholipid stereochemistry of the glycerol moiety has been directly determined for a surprisingly limited range of Bacteria and Archaea. Since the initial full structural characterisation of archaeol by Kates (1978), most subsequent studies of ether membrane lipids have assumed their stereochemistry while focusing on other aspects of their structure. Those studies that have determined the glycerol stereochemistry of membrane lipids (i.e. Sinninghe Damsté, et al. 2002a; Weijers et al. 2006) are largely consistent with the idea that it is a conserved difference between Bacteria and Archaea. Nonetheless, there is evidence that some Bacteria can make G1P-linked ether lipids. For example, the model bacterium *Bacillus subtilis* has been shown to possess homologues of archaeal G1PDH and GGGP (Guldan et al. 2008; Guldan et al. 2011). These enzymes allow *B. subtilis* to synthesise a typically archaeal ether link between G1P and HepPP, resulting in a lipid with archaeal characteristics, although there is no evidence that these archaeal-like lipids are used to make phospholipids, or are incorporated into the *B. subtilus* membrane.

Apart from stereochemistry, other characteristics of membrane phospholipids appear to be more variable, showing a mixture of archaeal and bacterial features. For example, the plasmalogens of animals and anaerobic Bacteria include an ether bond (Goldfine 2010).Branched glycerol dialkyl glycerol tetra-ether (brGDGT) lipids found in the environment have bacterial stereochemistry and branched rather than isoprenoidal alkyl chains, but they also contain ether bonds and span the membrane, as observed for canonical archaeal lipids (Schouten et al. 2000; Weijers et al. 2006). These brGDGTs are particularly abundant in peat bogs and were thought to be produced by Bacteria as adaptations to low pH environments (Weijers et al. 2006; Sinninghe Damsté et al. 2007), but are now known to occur in a wide range of soils and aquatic settings (Schouten et al. 2013). The enzymes responsible for their synthesis are currently unknown. On the other side of the “lipid divide”, some Archaea have been shown to produce membrane lipids with fatty acid chains and ester bonds (Gattinger et al. 2002). The biosynthetic pathways for all of these mixed-type membrane lipids remain unclear. However, given the frequency with which prokaryotes undergo horizontal gene transfer (Garcia-Vallvé et al. 2000), one possibility is that these mixed biochemical properties reflect biosynthetic pathways of mixed bacterial and archaeal origin.

A number of previous studies have investigated the evolutionary origins of phospholipid biosynthesis genes in Bacteria and Archaea using phylogenetic approaches, in order to test hypotheses about the nature of membranes in the earliest cellular life-forms (Peretó et al. 2004; Koga 2014; Villanueva et al. 2016; Yokobori et al. 2016). In this study, we build upon that work by performing comprehensive phylogenetic analyses for the core phospholipid biosynthesis genes in Bacteria and Archaea: the enzymes that establish membrane lipid stereochemistry and attach the two carbon chains to the glycerol phosphate backbone (Figure 1), as the histories of these enzymes are key to understanding the evolution of membrane biosynthesis and stereochemistry. Our analyses take advantage of the wealth of new genome data from environmental prokaryotes that has become available recently, and we employ new approaches for rooting single gene trees in order to circumvent some of the difficulties inherent in traditional outgroup rooting for anciently diverged genes. Our results agree with previous work in suggesting that LUCA likely possessed a cell membrane. Our rooted gene trees indicate that transfers of lipid biosynthetic genes from Archaea to Bacteria have occurred more frequently in evolution, particularly during the early diversification of the two domains.

## Results and Discussion

### Inter-domain horizontal transfer of core phospholipid biosynthesis genes

We performed BLASTp searches for the enzymes of the canonical archaeal and bacterial lipid biosynthesis pathways (Figure 1) against all archaeal and bacterial genomes in the NCBI nr database. Our BLAST searches revealed homologues for all of the core phospholipid biosynthesis genes of both pathways in both prokaryotic domains, with the exception of bacterial enzymes PlsB and PlsX, which we did not find in Archaea. Orthologues of the canonical archaeal genes are particularly widespread in many bacterial lineages (Table 1). Of the 48 bacterial phyla surveyed, 10 had no orthologues of the archaeal genes (Table 1, indicated by †). 6 phyla have orthologues of all three archaeal genes distributed across various genomes (Table 1, highlighted in yellow). Of these phyla, Firmicutes (genera *Bacillus* and *Halanaerobium*), Actinobacteria (genus *Streptomyces*) and Fibrobacteres (genera *Chitinispirillum* and *Chitinivibrio*) contain species which have all three genes in their genomes (Table 1, highlighted yellow with asterisk). Based on the presence of all three core biosynthetic genes, and given their recognised role in the synthesis of archaeal-like lipid components in *B. subtilis* (Guldan et al. 2008; Guldan et al. 2011), members of Firmicutes, Actinobacteria and Fibrobacteres lineages of Bacteria may be capable of making archaeal-like lipids, although we cannot determine if these are used in the production of membrane phospholipids. Of the 12 FCB group phyla we surveyed (Fibrobacteres, Chlorobi, Bacteroidetes and related lineages), all 12 have GGGPS and DGGGPS orthologues, but only Fibrobacteres have G1PDH orthologues (see Figure 1 for overview of pathway). In these species lacking G1PDH, it is unclear whether GGGPS and DGGGPS are active and if so, what they are used for; one possibility is that they catalyse the reverse reaction, catabolising archaeal lipids as an energy source. However, a very recent report (Villanueva et al. 2018) has shown that the GGGPS and DGGGPS genes from one FCB lineage, Cloacimonetes, support the production of archaeal-type membrane phospholipids and a mixed membrane when heterologously expressed in *E. coli*. This suggests that both *E. coli* and perhaps Cloacimonetes have an alternative, as yet unknown mechanism for making G1P, and that some FCB members may have mixed archaeal and bacterial membranes

Orthologues of the canonical bacterial genes are less widespread in Archaea (Table 1). Of all the genomes surveyed, none contained all homologues, although *Lokiarchaeum* sp. GC14_75 contained Glp, GlpK, PlsC and PlsY. Of the 20 phyla shown in Table 1, more than half (11) had no bacterial homologues in any of their genomes. Orthologues of GpsA, Gpl and Gpk are found in at least one genome of each of the major archaeal clades (Euryarchaeota, TACK, Asgardarchaeota and DPANN (Williams et al. 2017)). However, they appear sporadically. Within Euryarchaeota, of the seven classes surveyed, GpsA and Glpk appear in the genomes of four and Glp in five. Within the TACK superphylum, Glp and GlpK appear in Crenarchaeota and Korarchaeota, while GpsA appears only in a single crenarchaeote genome (*Thermofilum*).GpsA and Glp are also found in at least one genome in two of the 11 DPANN phyla surveyed (Woesearchaeota and GW2011, and Woesearchaeota and Parvarchaeota respectively), while GlpK is found in a single parvarchaeote genome (Candidatus Parvarchaeum acidiphilum ARMAN-4). Within the Asgardarchaeota superphylum, no orthologues for GpsA are found, and only one of the genomes (*Lokiarchaeum* sp. GC14_75) has Glp or GlpK. PlsC and PlsY are more restricted, being found mainly in environmental lineages within Euryarchaeota (Marine Groups II/III, all in class Thermoplasmatales), DPANN and Asgardarchaeota (Table 1).

### Early origins of archaeal-type membrane phospholipid biosynthesis genes in Bacteria

To investigate the evolutionary histories of membrane phospholipid biosynthesis, we inferred Bayesian single-gene phylogenies from the amino acid alignments using PhyloBayes 4.1 (Lartillot and Philippe 2004; Lartillot et al. 2007). We selected the best-fitting substitution model for each gene according to its Bayesian Information Criterion (BIC) score using the model selection tool in IQ-Tree (Nguyen et al. 2015). We used two complementary approaches to root these single-gene trees: a lognormal uncorrelated molecular clock in BEAST (Drummond and Rambaut 2007; Drummond et al. 2012), and the recently-described minimal ancestor deviation (MAD) rooting method of Tria et al. (2017). The MAD algorithm finds the root position that minimises pairwise evolutionary rate variation, averaged over all pairs of taxa in the tree. Many of our single gene trees were poorly resolved; since the existing implementation of the MAD algorithm (Tria et al. (2017)) does not explicitly incorporate topological uncertainty, we used MAD to root all of the trees sampled during the MCMC run, summarising posterior root support in the same way as for the molecular clock analyses; in the discussion below, we use the maximum *a posteriori* root as a point estimate for comparison between the two methods. For the genes for which an outgroup was available (G1PDH, GpsA and Glp, following Yokobori et al. 2016), we compared our results to traditional outgroup rooting. For more details, see Materials and Methods.

Glycerol-1-phosphate dehydrogenase (G1PDH) is the enzyme that establishes phospholipid stereochemistry in Archaea. Interestingly, the majority of the bacterial G1PDH orthologues do not appear to be recent horizontal acquisitions from Archaea, but instead form a deep-branching clan (Wilkinson et al. 2007) (PP = 1), resolved as sister to an archaeal lineage clan. The relationships within the clans are poorly resolved. The root position that receives the highest posterior support in the relaxed molecular clock analysis is that between the archaeal and bacterial clans, with a marginal posterior probability of 0.68 (Supplementary Table 1). This is substantially higher than the next most probable position, which places the root within the Bacteria with a posterior probability of 0.1. When rooted using MAD, the same root between the bacterial and archaeal clans is recovered with a marginal posterior probability of 0.62, also substantially higher than the next most probable root of 0.1. Rooting single genes trees can prove difficult, and this uncertainty is captured in the low root probabilities inferred using both the molecular clock and MAD methods. However, these analyses can be used to exclude the root from some regions of the trees with a degree of certainty. In the case of G1PDH, a post-LUCA origin of the gene would predict a root on the archaeal stem or within the archaea. In our analyses, no such root position has a significant probability (i.e. PP>0.05), and therefore the root is highly unlikely to be within the archaea. This is similar to topologies recovered by Peretó et al. (2014) and Carbone et al. (2015). The bacterial clan mainly comprises sequences from Firmicutes and Actinobacteria, with most of the other Bacteria grouping together in a single, maximally supported (PP = 1) lineage suggestive of recent horizontal acquisition from the Firmicutes/Actinobacteria clade, followed by further HGT.

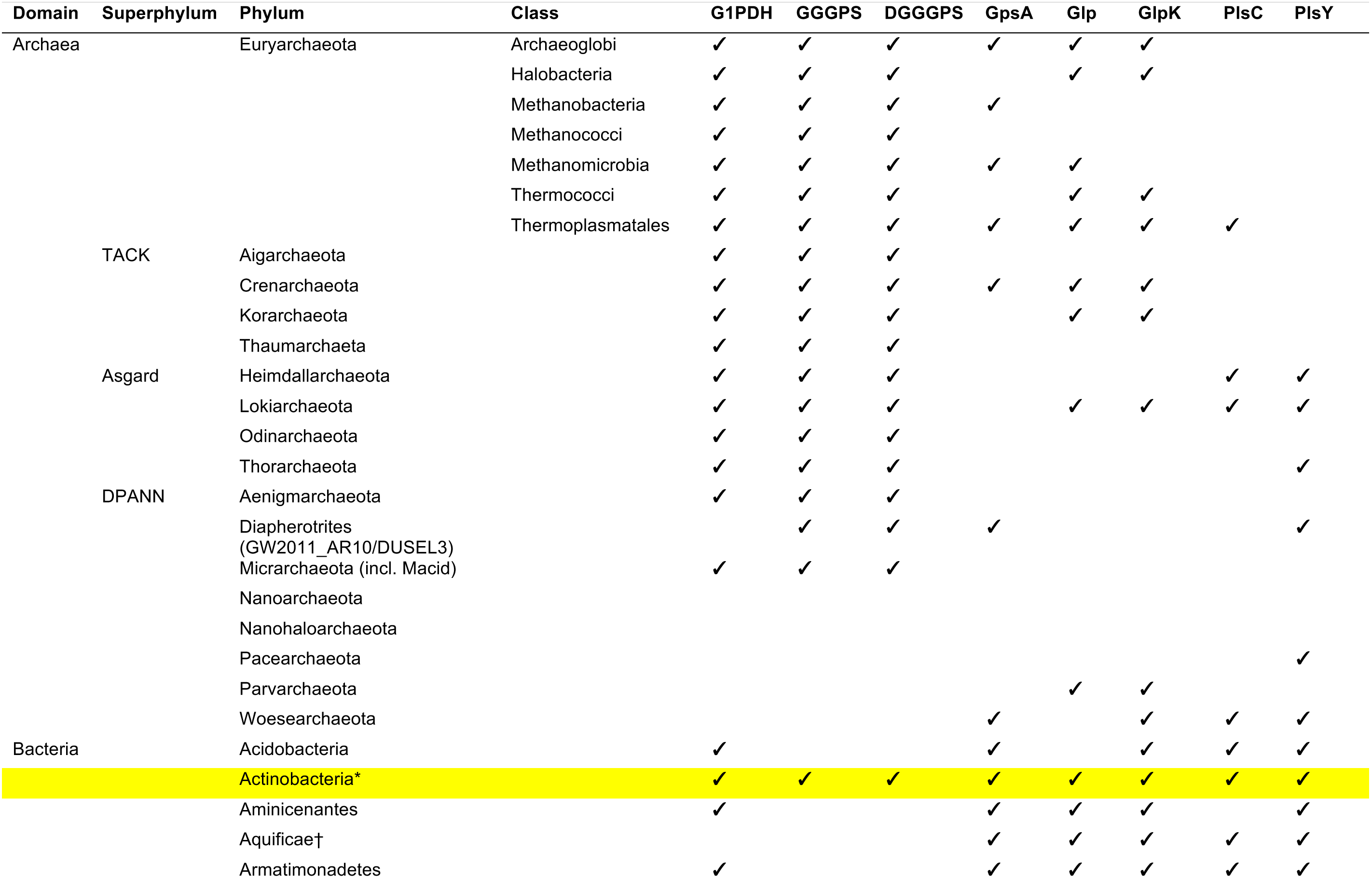
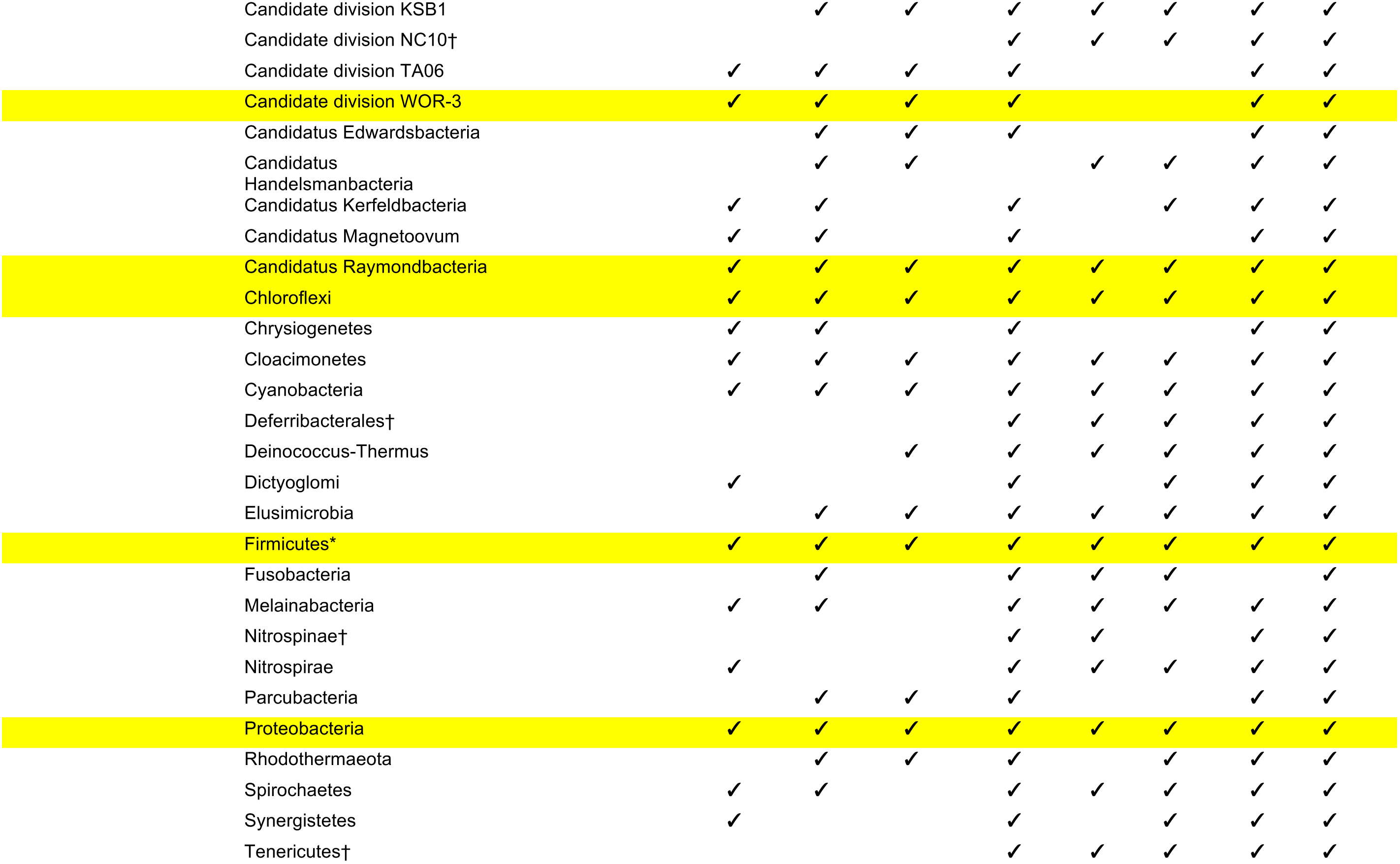
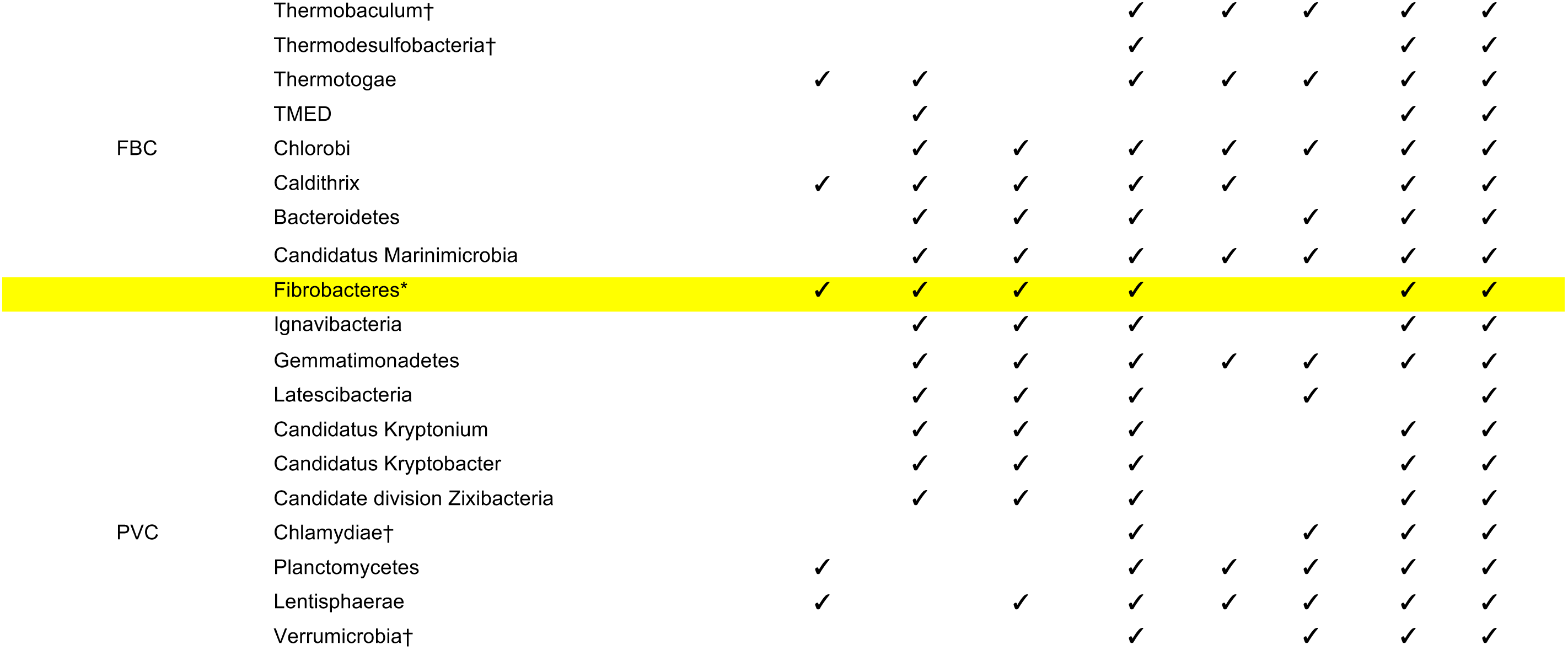

**Figure 2.**
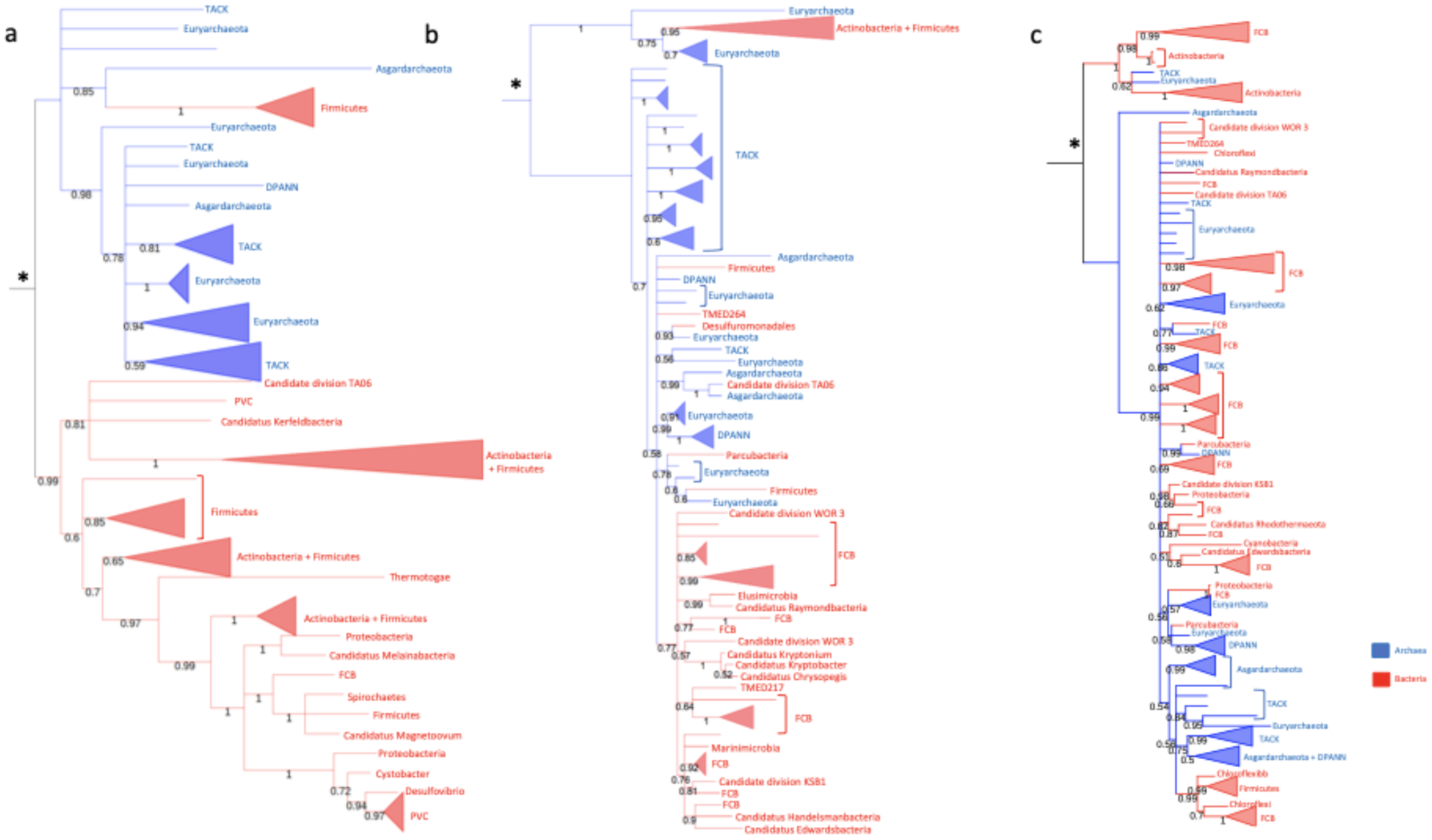
Bayesian consensus trees of archaeal enzymes, rooted using the uncorrelated lognormal molecular clock. Support values are Bayesian posterior probabilities, and the asterisk denotes the modal root position obtained using the MAD approach. Archaea in blue and Bacteria in red **a)** G1PDH tree (111 sequences, 190 positions) inferred under the best-fitting LG+C60 model. **b)** GGGPS tree (133 sequences, 129 positions) inferred under the best-fitting LG+C40 model. **c)** DGGGPS tree (97 sequences, 119 positions) inferred under the best-fitting LG+C60 model. FCB are Fibrobacteres, Chlorobi and Bacteroidetes, and related lineages. PVC are Planctomycetes, Verrucomicrobia and Chlamydiae, and related lineages. TACK are Thaumarchaeota, Aigarchaeota, Crenarchaeota and Korarchaeota. DPANN include Diapherotrites, Parvarchaeota, Aenigmarchaeota, Nanoarchaeota and Nanohaloarchaeota, as well as several other lineages. For full trees, see Supplementary Figures 1-4. For full unrooted trees see Supplementary Figures 16-18.

This root position is consistent with two scenarios that we cannot distinguish based on the available data. One possibility is an early transfer of G1PDH from stem Archaea into Bacteria, either into the bacterial stem lineage with subsequent loss in later lineages, or into the ancestor of Actinobacteria and Firmicutes, with subsequent transfers to other Bacteria. Alternatively, G1PDH could have already been present in LUCA, and was subsequently inherited vertically in both Archaea and Bacteria, followed by loss in later bacterial lineages. The Firmicute sequences within the archaeal clade appear to be a later transfer into those Firmicutes, apparently from Thorarchaeota.

Geranylgeranylglyceryl phosphate synthase (GGGPS) attaches the first isoprenoid chain to G1P. Phylogenetic analysis of GGGPS (Figure 2b) evidenced two deeply divergent paralogues, with the tree confidently rooted between them using both the relaxed molecular clock (PP = 0.99) and MAD methods (PP = 1) (Supplementary Table 1); resolution within each of the paralogues was poor. The recovery of two distinct paralogues has been noted in several previous studies (Nemoto et al. 2003; Boucher et al. 2004; Lombard et al. 2012b; Peterhoff et al. 2014). One of these paralogues comprises sequences from some Euryarchaeota (including members of the Haloarchaea, Methanomicrobia and Archaeoglobi), along with Firmicutes and Actinobacteria. The other paralogue comprises sequences from the rest of the Archaea – including other Euryarchaeota - and a monophyletic bacterial clade largely consisting of members of the FCB lineage. Taken with the root position between the two paralogues, the tree topology implies an ancestral duplication followed by sorting out of the paralogues and multiple transfers into Bacteria. Since genes from both GGGPS paralogous clades have been experimentally characterised as geranylgeranylglyceryl phosphate synthases (Nemoto et al. 2003; Boucher et al 2004), it appears that this activity was already present in LUCA before the radiation of the bacterial and archaeal domains. Payandeh (2006) has suggested, however, that the firmicute sequences (which comprise the majority of the sequences in the smaller paralogue) are used in teichoic acid synthesis. In this case, two apparently diverging paralogues may be an artifact due to changes in the sequences during neofunctionalisation. Lombard et al. (2012b), who also find two divergent homologues, and homologues in a large diversity of FCB bacteria (mostly Bacteroidetes), suggest that one of these homologues was likely present in the last archaeal common ancestor (LACA), whereas the bacterial sequences were likely horizontal transfers. To improve resolution among the deeper branches of the tree, we inferred an additional phylogeny focusing just on the larger of the two clades (Supplementary Figure 3). The root within this paralogous sub-tree fell between reciprocally monophyletic archaeal and bacterial clades (PP = 0.8, much higher than the next most likely root, within the bacteria, with PP = 0.07), suggesting that the gene duplication at the base of the GGGPS tree pre-dates LUCA.

Digeranylgeranylglyceryl phosphate synthase (DGGGPS) attaches the second isoprenoid chain to G1P. DGGGPS is present in all sampled Archaea, with the exception of three of the DPANN metagenome bins. Although the DGGGPS tree is poorly resolved (Figure 2c), both the molecular clock and MAD root the tree between two clades (PP = 0.43 and 0.79 respectively) (Supplementary Table 1) The smaller clade comprises mostly bacterial sequences form the Actinobacteria and FCB lineages, as well as two archaeal archaeal sequences (from the TACK and Euryarchaeota lineages). The larger clade contains sequences form a diversity of Bacteria, particularly FCB (also reported by Villanueva et al. 2018), as well as Archaea.DGGGPS is part of the UbiA protein superfamily, which are involved in a number of different biosynthetic pathways, including the production of photosynthetic pigments, and are therefore widely distributed in Bacteria, and are known to have undergone extensive HGT (Hemmi 2004). Indeed, several of the sequences used in our analyses (and those in previous studies, such as Villanueva et al. 2016) are annotated on NCBI as other proteins within this superfamily (see Supplementary Table 3). To distinguish orthologues of DGGGPS from other, distantly related members of the UbiA superfamily that might have different functions, we inferred an expanded phylogeny including our initial sequence set and sequences sampled from the other known UbiA sub-families (Supplementary Figure 25). Surprisingly, this analysis indicated that the Thaumarchaeota lack an orthologue of the DGGGPS gene that other Archaea use to attach the second isoprenoid chain; the most closely related Thaumarchaeota sequences branch within another UbiA sub-family with high posterior support (PP = 0.99). Thaumarchaeota may be using this paralogue to perform the same function, or may use another unrelated enzyme to catalyze this reaction. The wide distribution of this enzyme across both Archaea and Bacteria, and the occurrence of both domains on either side of the root, for both rooting methods, suggest either multiple transfers into Bacteria from Archaea, or that DGGGPS was present in LUCA and inherited in various archaeal and bacterial lineages, followed by many later losses in and transfers between various lineages.

In sum, our results of archaeal phospholipid biosynthesis genes suggest that there have been repeated, independent inter domain transfers of these genes from Archaea to Bacteria throughout the evolutionary history of life. Furthermore, our phylogenetic analyses do not exclude the possibility that the genes of the archaeal pathway were present in LUCA. If correct, this would imply that LUCA had the capability to make archaeal-type membrane phospholipids.

### Transfers of bacterial membrane phospholipid genes into Archaea

In contrast to our analyses of proteins of the classical archaeal pathway, phylogenies of proteins of bacterial-type membrane phospholipid biosynthesis pathways suggested that their orthologues in Archaea were the result of relatively recent horizontal acquisitions. Homologues of both forms of G3PDH and GlpK are broadly distributed in Archaea, however these three enzymes are not exclusive to phospholipid synthesis, and have been shown to be used in glycerol metabolism in some autotrophic Archaea (Nishihara et al. 1999). Of the enzymes thought to function exclusively in bacterial membrane phospholipid biosynthesis, we did not find any archaeal homologues for PlsB or PlsX. Archaeal PlsC and PlsY homologues are patchily distributed, and are found only in metagenomic bins. It therefore seems unlikely that any of these genes function in membrane phospholipid synthesis in Archaea.

The root positions for each of the trees using both molecular clock and MAD have low posterior probabilities (Supplementary Table 1), so that the exact root positions are unclear; however, we did not observe any support for root positions outside the Bacteria. This is consistent with the hypothesis that the core bacterial pathway first evolved after the bacterial lineage diverged from LUCA. *GpsA* and *glp* are two genes that code for glycerol-3-phosphate (G3PDH), which establishes phospholipid stereochemistry in Bacteria. The deep relationships between the archaeal and bacterial sequences in the GpsA tree are poorly resolved (Figure 3a), while being better resolved for Glp (Figure 3b). The root position in both trees is poorly resolved for both rooting methods (Supplementary Table 1). The highest marginal posterior probability for the root positions recovered in the GpsA tree are 0.31 and 0.59 and for the molecular clock and MAD respectively, and 0.5 and 0.44 respectively for Glp. The tree inferred for GlpK --- the gene that codes for glycerol synthase, which can synthesise G3P from glycerol (Figure 4a)---shows a similar pattern to the phylogenies of GpsA and Glp. Again, the root positions have low posterior support (0.47 and 0.34 for the molecular clock and MAD respectively). However, in each case, there is evidence of recent transfers from Bacteria to Archaea, as we recover several distinct bacterial and archaeal clades with moderate to high support (0.8-1), as also reported by Villanueva et al. (2016).

**Figure 3.**
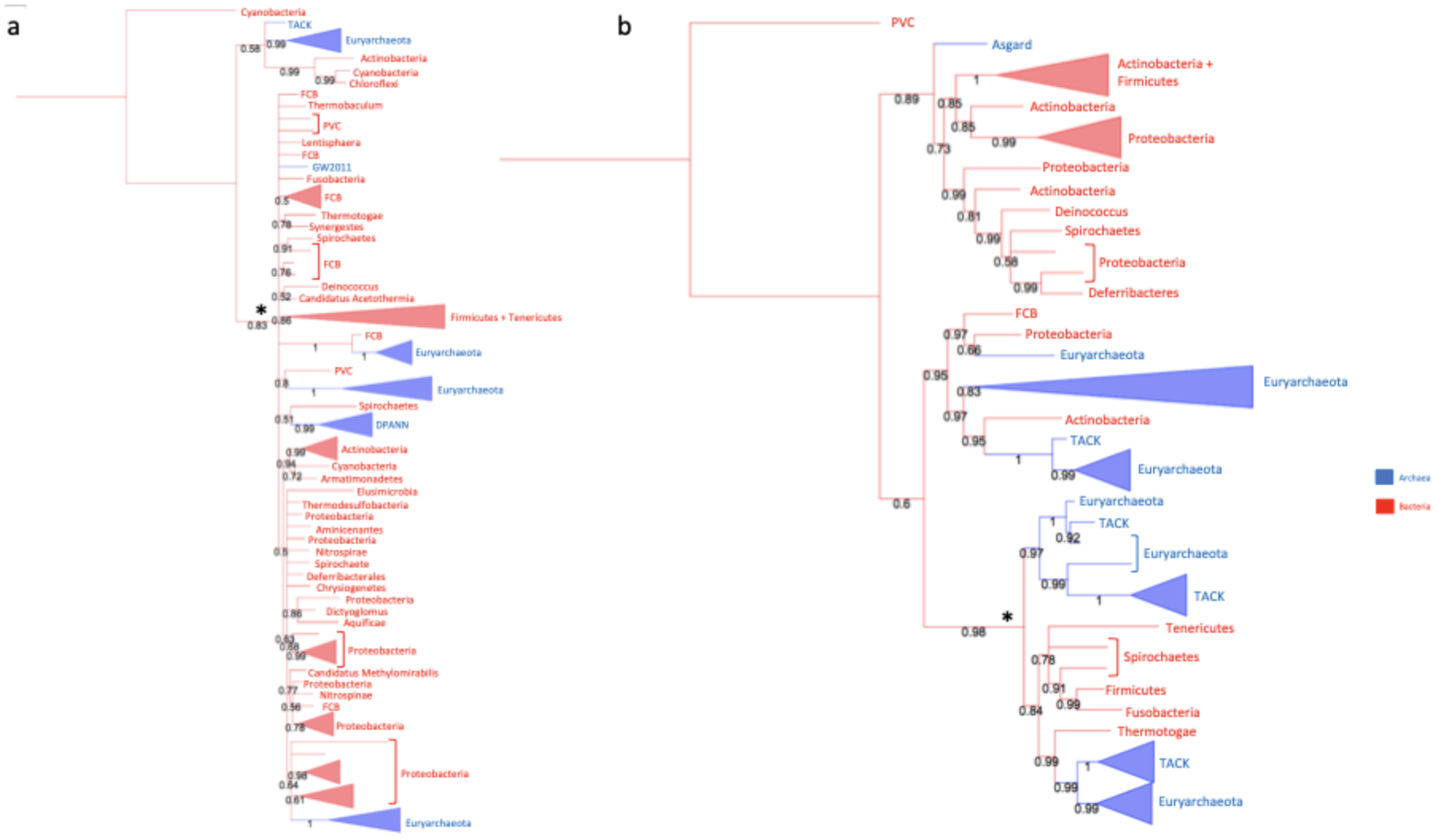
Bayesian consensus trees of both G3PDH enzymes, rooted using the uncorrelated lognormal molecular clock. Support values are Bayesian posterior probabilities, and the asterisk denotes the modal root position obtained using the MAD approach. Archaea in blue and Bacteria in red. **a)** gpsA tree (84 sequences, 169 positions) inferred under the best-fitting LG+C60 model **b)** glp tree (51 sequences, 199 positions) inferred under the best-fitting LG+C40 model. FCB are Fibrobacteres, Chlorobi and Bacteroidetes and related lineages. PVC are Planctomycetes, Verrucomicrobia and Chlamydiae and related lineages. TACK are Thaumarchaeota, Aigarchaeota, Crenarchaeota and Korarchaeota. DPANN include Diapherotrites, Parvarchaeota, Aenigmarchaeota, Nanoarchaeota and Nanohaloarchaeota, as well as several other lineages. For full trees, see Supplementary Figures 5-6. For full unrooted trees see Supplementary Figures 19-20.

PlsC and PlsY (which attach fatty acids to G3P) both have many fewer orthologues among archaeal genomes, all of which are derived from environmental samples (Embley and Martin 2006; Martin et al. 2015; Eme et al. 2017). Both trees are poorly resolved (Figure 4b). Both are rooted within the Bacteria, with PlsC having the low posterior of 0.28 (with the next most likely, also within the Bacteria, being 0.1). The PlsY (Fig 4c) has a more certain root position, with a posterior of 0.57, and the next most probable being 0.1. For PlsY, MAD recovers the same root as the molecular clock, with a high posterior probability (0.85). When the PlsC tree is rooted using MAD, the root is resolved between two clades, which are not recovered in the inferred tree topology and has a low posterior probability of 0.03. All of the archaeal homologues seem to horizontal acquisitions from Bacteria.

**Figure 4.**
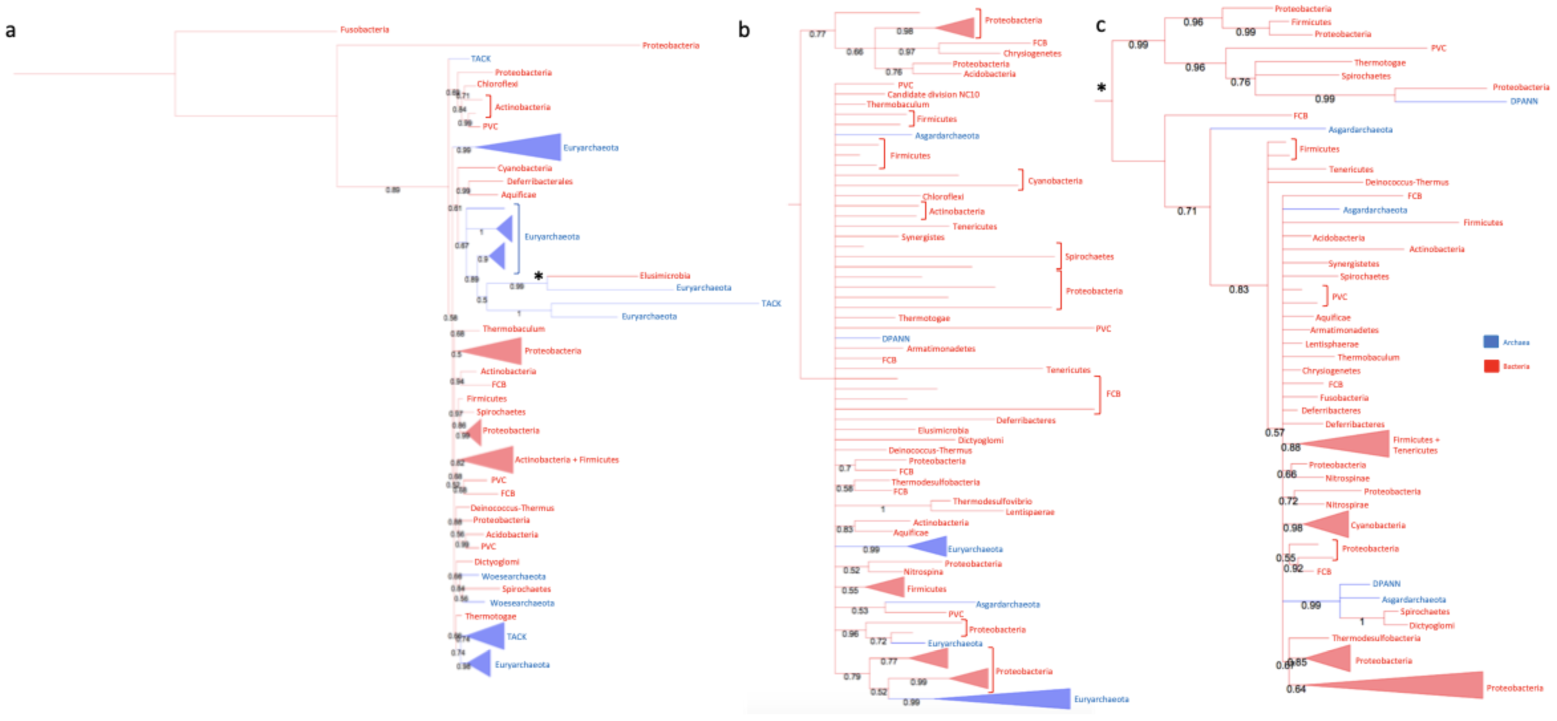
Bayesian consensus trees of glpK, PlsC and PlsY enzymes, rooted using the uncorrelated lognormal molecular clock. Support values are Bayesian posterior probabilities, and the asterisk denotes the modal root position obtained using the MAD approach. Archaea in blue and Bacteria in red. **a)** glpK tree (77 sequences, 363 positions) inferred under the best-fitting LG+C60 model **b)** PlsC tree (74 sequences, 57 positions) inferred under the best-fitting LG+C50 model **c)** PlsY tree (60 sequences, 104 positions) inferred under the best-fitting LG+C60 model. FCB are Fibrobacteres, Chlorobi and Bacteroidetes and related lineages. PVC are Planctomycetes, Verrucomicrobia and Chlamydiae and related lineages. TACK are Thaumarchaeota, Aigarchaeota, Crenarchaeota and Korarchaeota. DPANN include Diapherotrites, Parvarchaeota, Aenigmarchaeota, Nanoarchaeota and Nanohaloarchaeota, as well as several other lineages. For full trees, see Supplementary Figures 7-9. For full unrooted trees see Supplementary Figures 20-23.

### Comparing outgroup and outgroup-free rooting for single gene trees

Evolutionary interpretations typically depend on rooted trees, but rooting single gene trees can prove difficult. The most widely-used approach is to place the root on the the branch leading to a predefined outgroup (Penny 1976). However, this can be challenging for ancient genes when closely-related outgroups are lacking; either the outgroup method cannot be used at all, or else the long branch leading to the outgroup can induce errors in the in-group topology (a phenomenon known as long branch attraction (LBA); see Gouy et al. 2015).

In the case of phospholipid biosynthesis, some of the key genes belong to larger protein families whose other members, although distantly related, have conserved structures and related functions (Peretó et al. 2004). Several previous studies looking at the history of phospholipid biosynthetic genes have used these outgroups for rooting. Due in part to the difficulties of outgroup rooting for ancient genes, these studies have disagreed on the roots for some of these gene trees, leading to very different evolutionary conclusions. Our outgroup-free results are consistent with those of Peretó et al. (2004) and Carbone et al. (2015), but not with those of the recent study of Yokobori et al. (2016). Yokobori et al. used outgroups to root trees for G1PDH, G3PDH (both GpsA and Glp) and GlpK. Their root inferences differed from ours in that they found that bacterial G1PDH sequences formed a monophyletic group that branched from within Archaea, suggesting more recent horizontal transfer from Archaea to Bacteria, as opposed to transfer from stem Archaea or vertical inheritance from LUCA (Fig. 2a). On the other hand, their analysis of Glp recovered Bacteria on one side of the root, and a clade of Bacteria and Archaea on the other. They interpreted this as evidence for the presence of Glp in LUCA, and therefore that LUCA would have had bacterial-type G3P membrane phospholipids.

Single-matrix models, such as those used by Yokobori et al. (2016), have been shown to be more susceptible to phylogenetic artifacts such as LBA than the profile mixture models used here (Lartillot et al. 2007). To investigate whether the differences in root inference between our analyses and those of Yokobori et al. (2016) might be the result of LBA, we performed outgroup rooting analysis on G1PDH, GpsA and Glp, augmenting our datasets with a subsample of the outgroups used by Yokobori et al. and using the same models used to infer the unrooted trees (LG+C60 in each case). The resulting trees (Supplementary Figures 10-12) show different topologies when compared to the unrooted trees (Supplementary Figures 16, 19-20). This suggests that the long branch outgroup may be distorting the ingroup topology.

We also performed model testing in IQ-Tree and compared the fit of the chosen models to the models used by Yokobori et al. (see Material and Methods below). LG+C60 was selected for both G1PDH and Glp, while LG+C50 was selected for Gpsa (Supplementary figure 24). The results of these analyses indicate that the empirical profile mixture models which we have used here fits each of these alignments significantly better than the single-matrix models of Yokobori et al. (Supplementary Table 2). However, even analyses under the best-fitting available models show distortion of the ingroup topology upon addition of the outgroup (Supplementary Figures 10-12, 24), when compared to the unrooted topologies (Supplementary Figures 16, 19-20). In each case, we found the root in a different place to those recovered by Yokobori et al. In the G1PDH tree, we find Bacteria (Firmicutes) to be most basal, rather the Crenarchaeota found by Yokobori. In the case of GpsA, Yokobori et al. did not find compelling support for an origin in LUCA, but they did recover one archaeal lineage (the Euryarchaeota) at the base of the in-group tree with low (bootstrap 48) support. While our GpsA tree is also poorly resolved, we do not find evidence to support the basal position of the archaeal lineages, and therefore for the presence of GpsA in LUCA. For glp, which Yokobori et al. trace back to LUCA due to the basal position of the archaeal sequences, the outgroup sequences did not form a monophyletic group, and were instead distributed throughout the tree (Supplementary Figure 11). Thus, analyses under the best-fitting available models did not support the presence of bacterial lipid biosynthesis genes in LUCA. Further, the distortion of the ingroup topologies suggests that these outgroups may not be suitable for root inference, at least given current data and methods. The relaxed molecular clock and the MAD method have their own assumptions and limitations, but these results suggest that they may be useful for rooting trees in other contexts, either as part of a sensitivity test or when suitable outgroups are not available.

### Origin of eukaryotic membrane phospholipid biosynthesis genes

Phylogenetics and comparative genomics suggest that eukaryotes arose from a symbiosis between an archaeal host cell and a bacterial endosymbiont that evolved into the mitochondrion (reviewed, from a variety of perspectives, in Embley and Martin 2006; Martin et al. 2015; Eme et al. 2017; Roger et al. 2017). Genomic and phylogenetic evidence indicates that the host lineage belonged to the Asgardarchaeota superphylum (Spang et al. 2015; Zaremba-Niedzwiezka et al. 2017). The origin of bacterial-type membrane phospholipids in eukaryotes is therefore an important evolutionary question that has received considerable attention (Woese et al. 1990; Kandler 1995; López-Garcia and Moreira 2006; Baum and Baum 2014; Gould et al. 2016). Given the evidence for transfer of bacterial-type phospholipid biosynthesis genes into Archaea, one possibility - also raised by the results of Villanueva et al. (2016) - is that eukaryotes may have inherited their bacterial lipids vertically from the archaeal host cell. Both our study and that of Villanueva et al. (2016) point to the presence of orthologues for bacterial lipid genes in Asgardarchaeota. These include Glp, PlsC and PlsY orthologues in *Lokiarchaeum* sp. GC14_75, PlsC and PlsY in Heimdallarchaeota archaeon LC_2, and PlsY in Thorarchaeota archaeon SMTZ1-83 (Table 1). However, phylogenies of these genes (Supplementary Figures 13-15) do not support a specific relationship between eukaryotes and any of the archaeal sequences, and so do not provide any compelling support for an origin of eukaryotic lipids via the archaeal host cell.

### Conclusions

Our phylogenetic analyses of lipid biosynthesis genes support two main conclusions about prokaryotic cell physiology and early cell evolution. First, our results corroborate previous evidence for extensive horizontal transfer of lipid genes, particularly from Archaea to Bacteria, from potentially very early to more recent evolutionary times. The functions of these genes remain unclear, but in *B. subtilis* (Guldan et al. 2008; Guldan et al. 2011) they are involved in making archaeal-type G1P ether-linked lipids, while in the FCB lineage Cloacimonetes (Villanueva et al. 2018) they may be involved in synthesising archaeal-type phospholipids that are incorporated into the bacterial cell membrane. Evidence that these genes have undergone horizontal transfer, both early in evolution and more recently, provides a potential mechanism for the remarkable diversity of membrane lipids, and especially ether lipids, in environmental settings (Schouten et al. 2001). We also note that it is intriguing that bacterial lipids with archaeal features are particularly abundant in settings characterised by high archaeal abundances, including cold seeps, wetlands and geothermal settings (Schouten et al. 2013), potentially providing ecological opportunity for gene transfer. Experimental work to characterise the enzymes that make these environmental lipids will be needed to test this prediction.

A second, and more tentative, result of our study relates to the antiquity of the canonical archaeal and bacterial pathways. Our analyses suggest that the enzymes for making G1P lipids may have been present in the common ancestor of Archaea and Bacteria. Under the consensus view that the root of the tree of life lies between Bacteria and Archaea, this would imply that LUCA could have made archaeal type membranes. This finding is intriguing in light of previous work suggesting the presence of isoprenoids produced by the mevalonate pathway in LUCA (Lombard and Moreira 2011; Castelle and Banfield 2018). By contrast, the roots for the bacterial genes were weakly resolved within the bacterial domain. There is therefore no positive evidence from our trees to suggest that the bacterial pathway was present in LUCA, although we cannot exclude this possibility. The consensus universal root between Bacteria and Archaea is supported by analyses of ancient gene duplications (Gogarten et al. 1989; Iwabe et al. 1989; Zhaxybayeva et al. 2005) and genome networks (Dagan et al. 2010), but some analyses have supported an alternative placement of the root within Bacteria (Cavalier-Smith 2006; Lake et al. 2009; Williams et al. 2015). Our trees do not exclude a within-Bacteria root, in which case LUCA would have possessed the bacterial pathway, and the archaeal pathway would have evolved along the archaeal stem, or in a common ancestor of Archaea and Firmicutes (Cavalier-Smith 2006; Lake et al. 2009).

If one membrane lipid pathway evolved before the other, this would imply that one of the two prokaryotic lineages changed its membrane lipid composition during early evolution. The evolutionary processes that drive such changes remain unclear, in part because we still do not fully understand the functional differences between modern archaeal and bacterial membranes. Compared to bacterial-type membranes, archaeal-type membranes maintain their physiochemical properties over a broader range of temperatures, and may be more robust to other environmental extremes (Van de Vossenberg et al. 1998; Koga 2012). If the archaeal pathway is older than the bacterial pathway, then that could reflect a LUCA adapted to such extreme settings. It is then intriguing to speculate on the evolutionary drivers for subsequent adoption of bacterial-type membranes, especially since the Bacteria appear to be more successful than the Archaea in terms of abundance and genetic diversity (Danovaro et al. 2016; Hug et al. 2016; Castelle and Banfield 2018). Moreover, an analogous change has happened at least once in evolutionary history, during the origin of eukaryotic cells (Martin et al. 2015). Chemical considerations suggest such bonds ought to be energetically cheaper to make and break, although we know of no published experimental data on these relative biosynthetic costs. Alternatively, bacterial-type membrane lipids comprise a variety of fatty acyl moieties, varying in chain length, unsaturation, degree of branching and cyclisation, and these could impart a degree of flexibility and adaptability that provides a marginal benefit in dynamic mesophilic environments. If so, that advantage could translate to bacterial ether lipids that are also widespread in non-extreme settings and also characterised by a variety of alkyl forms (Pancost et al. 2001). Conversely, if bacterial-type membranes were ancestral, the transition to archaeal-type membranes could have been driven by adaptation to high environmental temperatures: ether bonds are more thermostable than esters (Van de Vossenberg et al. 1998; Koga 2012), and are also found in the membranes of thermophilic Bacteria (Kaur et al. 2015). In any case, the widespread occurrence of bacterial-type, archaeal-type and mixed-type membrane lipids in a range of environments, as well as the widespread occurrence of the associated biosynthetic genes across both domains, suggests that except for high temperature and low pH settings, the advantages of either membrane type is marginal.

## Materials and Methods

### Sequence selection

For Archaea, we selected 43 archaeal genomes, sampled evenly across the archaeal tree. We took corresponding archaeal G1PDH, GGGPS and DGGGPS amino acid sequences from the data set of Villanueva et al. (2016) and performed BLASTp searches the find these sequences in genomes not included in that dataset. For Bacteria, we selected 64 bacterial genomes, sampled so as to represent the known genomic diversity of bacterial phyla (Hug et al. 2016). We used G3PDH *gpsa*, G3PDH *glp* and GlpK sequences from Yokobori et al. (2016) and performed BLASTp searches to find those sequences in bacterial species not in their data set. For PlsC and PlsY, we took the corresponding sequences form Villanueva et al. 2016, and performed BLASTp searches to find these sequences in the remaining genomes. For PlsB and PlsX, we searched for the respective terms in the gene database on the NCBI website, and upon finding well-verified occurrences, performed BLASTp searches to find the corresponding amino acid sequences in the remaining genomes. We then used BLASTp to look for bacterial orthologues of the archaeal enzymes and vice versa. We selected sequences that had an E-value of less the e-7 and at least 50% coverage. Accession numbers for sequences used are provided in Supplementary Table 3.

### Phylogenetics

The sequences were aligned in mafft (Katoh et al. 2002) using the --auto option and trimmed in BMGE (Criscuolo and Gribaldo 2010) using the BLOSUM30 model, which is most suitable for anciently-diverged genes. To construct gene trees from our amino acid sequences, we first selected the best-fitting substitution model for each gene according to its BIC score using the model selection tool in IQ-Tree (Nguyen et al. 2015). For all of the genes we analysed, the best-fitting model was a mixture model combining the LG exchangeability matrix (Le and Gascuel 2008) with site-specific composition profiles (the C40, C50 and C60 models (Lartillot and Philippe 2004; Le et al. 2008)) to accommodate across-site variation in the substitution process. LG+C60 was used for G1PDH, DGGGP, GpsA, Glp, GlpK and PlsC. LG+50 was used for PlsY. LG+C40 was used for GGGPS. A discretised Gamma distribution (Yang 1994) with 4 rate categories was used to model across-site rate variation. The trees were run with their respective models in PhyloBayes (Lartillot and Philippe 2004, 2006; Lartillot et al. 2007); convergence was assessed using the bpcomp and tracecomp programs (maxdiff < 0.1; effective sample sizes > 100), as recommended by the authors. We additionally ran all trees using the LG+C60 model (Supplementary Figures 26-29).

The trees were rooted with a lognormal uncorrelated molecular clock, using the LG model with a discretised Gamma distribution (Yang 1994) with 4 rate categories, and a Yule tree prior (Stadler 2009; Hartmann et al. 2010) in BEAST (Drummond 2007; Drummond et al. 2012). We also rooted the trees using minimal ancestral deviation (MAD) rooting (Tria et al 2017). MAD rooting requires a fully-resolved, bifurcating tree; since some parts of the consensus phylogenies were poorly resolved, we integrated over this topological uncertainty by computing the optimum MAD root position for each tree sampled during the MCMC analysis, and obtained marginal posterior probabilities for these root positions using RootAnnotator (Calvignac-Spencer et al. 2014).

For G1PDH, GpsA and Glp, we also rooted the trees using a subsample of the outgroup sequences used by Yokobori et al. (2016). The outgroups used were two 3-dehydroquinate synthase (DHQS), five glycerol dehydrogenase (GDH) and five alcohol dehydrogenase (ALDH) sequences for G1PDH; six hydroxyacyl-CoA dehydrogenase (HACDH) and six UDP-glucose 6-dehydrogenase (UDPGDH) sequences for GpsA; and 12 FAD-dependent oxidoreductase sequences for Glp. All three of these trees were inferred under the LG+C60 model to directly compare to the unrooted trees. Trees were also inferred from best fit models selected in IQTree (LG+C60 for G1PDH and Glp, and LG+C50 for GpsA).

Eukaryotic orthologues of prokaryotic phospholipid biosynthesis genes (Glp, GpsA and PlsC) were identified by performing BLASTp searches on 35 eukaryotic genomes from across eukaryotic diversity using *Homo sapiens* query as the sequence in each case, selecting sequences with an E-value of e-7 or less, and at least 50% coverage. We then performed model testing in IQTree and inferred trees in PhyloBayes using the selected substitution model (LG+C60 for PlsC and LG+C50 for Glp and GpsA).

All sequences, alignments and trees referred to in this study can be obtained from 10.6084/m9.figshare.6210137.

## Acknowledgements

GAC, RDP, and TAW conceived the project. GAC and TAW designed and performed the analyses. GAC, RDP and TAW interpreted the results and wrote the manuscript. GAC is supported by a Royal Society Research Grant to TAW. TAW is supported by a Royal Society University Research Fellowship. We thank George S. Attard, Anja Spang, T. Martin Embley and two anonymous reviewers for helpful feedback on earlier versions of the manuscript.

